# Gaussian Process Based Heteroscedastic Noise Modeling for Tumor Mutation Burden Prediction from Whole Slide Images

**DOI:** 10.1101/554261

**Authors:** Sunho Park, Hongming Xu, Tae Hyun Hwang

## Abstract

Tumor mutation burden (TMB) is a quantitative measurement of how many mutations present in tumor cells from a patient tumor as assessed by next-generation sequencing (NGS) technology. High TMB is used as a predictive biomarker to select patients that likely respond to immunotherapy in many cancer types; thus it is critical to accurately measure TMB for guiding patients to immunotherapy treatments.be used to predict genetic Recent studies showed that image features from histopathology whole slide images could be used to predict genetic features (e.g., mutation status) or clinical outcome of cancer patients. In this study, we develop a computational algorithm to predict the TMB level from cancer patients’ histopathology whole slide images. We formulate TMP prediction problem based on whole slide images as a multiple instance learning (MIL) problem. A whole slide image (a bag) is divided into multiple small image blocks/patches (instances), but a single label (e.g., TMB level) is available only to an entire whole slide image not to each image block. In particular, we propose a novel heteroscedastic noise model for MIL based on the framework of Gaussian process (GP), where the noise variance is assumed to be a latent function of image level features. This noise variance can encode the confidence in predicting the TMB level from each training image and make the method to put different levels of effort to classify images according to how difficult each image can be correctly classified. The proposed method tries to fit an easier image well while it does not put much effort into classifying a harder (ambiguous) image correctly for TMP prediction. Expectation and propagation (EP) is employed to infer our model efficiently and to find the optimal hyper-parameters. In experiments using the whole slide images from synthetic and real-world data sets from The Cancer Genome Atlas (TCGA), we demonstrate that our method outperforms base-line methods for TMP prediction including a special case of our method that does not include the heteroscedastic noise modeling and a multiple instance ordinal regression (MIOR) to solve ordinal regression in the MIL setting.

## 1 Introduction

Tumor mutation burden (TMB) measures the number of mutations found in a patient’s tumor assessed by next-generation sequencing (NGS) technology. Recent studies have shown that TMB is associated with better response to immunotherapy in many cancer types including lung and bladder cancer, thus can be used as a predictive biomarker for immunotherapy (*3, 10*). One key challenge is that not all patients would have adequate tumor tissue or will be able to undergo a tumor biopsy to generate NGS-based genomic testing to measure TMB. Although a blood-based TMB (e.g., liquid biopsies) recently becomes available, this approach also poises many technical challenges to measure TMB accurately. On the other hand, histopathology whole slide images used by pathologists to diagnose and assess tumor stage for cancer patients are widely available for the most cancer patients. Recent studies have shown that computational histopathology whole slide images (hereafter whole slide images) could be used to predict tumor subtype, clinical outcome, or certain mutation status (*6, 25, 31*). The fundamental assumption of these studies is that the morphology present in whole slides images is reflecting genetic aberrations driving cancer; therefore the algorithm utilizing morphological image features can reversely predict genetic status, clinical outcome, etc.

Our central hypothesize of this study is that the morphological image features generated from whole slide images could be used to predict TMB status. However, there are several difficulties to implement efficient TMB prediction methods from whole slide images. The main challenge is that the relation between TMB and morphology on images has not been fully explored. There has been no research to address this relation clearly yet. The second challenge is related to computational feasibility. Whole slide images are typically very huge: the size of either dimension of images is typically larger than 100,000 pixels and the file size of an image can be from hundreds of megabytes to several gigabytes. One possible solution to the computational difficulty is to divide an image into multiple non-overlapping image blocks. Then predictive or analysis methods are separately applied to each image block in an image, and then the final results can be obtained by aggregating the results from all the image blocks (*6, 25*). On the other hands, in (*31*) feature vectors from all the image blocks in an image are first combined into a single feature vector which is then fed into a SVM classifier as the input with the image’s label.

There has been few research to predict genetic features from histopathology images. Most research is designed to predict presence of certain mutations, e.g., EGFR in lung cancer (*6*), from images. As mentioned earlier, due to computational issues, the methods divide an image into multiple blocks and train predictive models on the images blocks (the target label of an image block is assumed to be the same as that of the image to which the block belongs) (*6, 25*). We can consider the same approach to predict TMB from whole side images. However, although this approach can increase the number of labeled training samples, it may degrade the predictive performance of the algorithms because image blocks that might be irrelevant to prediction are also treated as individual training samples and directly affecting the decision boundary. In fact, TMB prediction can be formulated as multiple instance learning (MIL) in which a label is assigned to a set (bag) of instances, not to each instance. In our problem, an image can be considered a bag and image blocks in an image instances. Furthermore the aforementioned methods, where all the image blocks in an image are assumed to have the same label as the image, corresponds to the naive MIL method. It is known that methods that fully account the MIL assumptions typically outperform the naive MIL method.

We develop a prediction method of TMB from whole slide images, where a novel noise model for MIL is proposed based on the framework of Gaussian process (GP). We formulate the prediction problem of TMB as ordinal regression (i.e., predicting TMB low ≾ intermediate ≾ high) and assume a latent function mapping from the input feature space to the intervals on the real line, modeled by a GP. Then, the mean value of the function values evaluated at all image blocks in an image is compared to the corresponding label in a noise (likelihood) model where the noise variance is also modeled by another GP (heteroscedastic noisy modeling) based on image level features that are calculated from feature vectors in an image. This noise variance encodes the confidence that our method has regarding image samples. The method more relies on image samples that are easier (having the higher confidence level or the lower noise variance) but less relies on image samples that are hard (having the higher noise variance). In other words, the method ties to fit easier image samples well while it does not put much effort in classifying harder (ambiguous) images correctly. Finally, expectation propagation (EP) is employed to efficiently train our model and find the optimal hyper-parameters. We have demonstrated from a synthetic and real-world data sets the usefulness of our heteroscedastic noise modeling. Our contribution can be summarized as follows

- To our best knowledge our work is the first attempt to solve the prediction of TMB from whole slide images in the framework of multiple instance learning (MIL).
- We propose a heteroscedastic noise model for MIL, which can encode the confidence in predicting TMB from an image.

## 2 Related Work

MIL, first introduced in (*7*), is a variation of supervised learning where a (positive or negative) label is assigned not to an individual instance but to a bag that consists of a set of instances. The standard MIL assumption (*7, 28*) states that a bag is positive if and only if there is at least one positive instance in the bag while a negative bag should not contain any positive instances. This assumption can be implemented by maximum operator (*1, 17, 19*). Assuming that each instance in the *i*th bag has a latent label *y_ij_* ∈ {0,1}, the standard MIL assumption states that the bag’s label can be given by

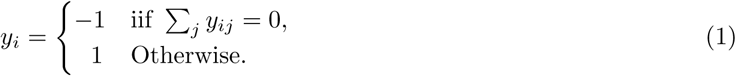

Furthermore, it can be further simplified using maximum operator as mentioned earlier, i.e., *y_i_* = max_*j*_{y_ij_}. In the standard MIL assumption, we can easily see that only instances in a bag with positive label can affect the class label of the bag. On the other hand, there have been several alternatives to the standard MIL assumption (please refer (*8*)). One of them is the collective assumption (*8, 32*), where all instances in a bag are assumed to equally contribute the bag’s label. Our prediction method is based on the collective assumption to connect image blocks (instances) in a patient’s whole slide image (a bag) to the patient’s TMB level, which will be explained in the next section.

Most of MIL algorithms are designed to solve binary classification problem (*11, 18, 19*). There have been proposed very few MIL methods for ordinal regression problems (*29*) as it is not straightforward to directly extend the standard MIL assumption to multi-class classification or ordinal regression problems. On the other hand, Gaussian process is a powerful Bayesian non-parametric method for nonlinear function estimation problem (*23*). It has been successfully applied to ordinal regression problems as well (*5*). In this study, we propose an efficient ordinal regression method for MIL based on the framework of GP. Compared to the method in (*29*), which gives point estimate solutions, our method could be more attractive because our method is a Bayesian approach, more robust to overfitting, and provides a way to find optimal hyper-parameters by maximizing marginal likelihood (or evidence).

Recently, deep learning based methods have been extensively applied to analyze whole slide images (*6, 16, 25*). These methods could be more desirable than traditional machine learning methods because the performance of traditional machine learning methods rely on the quality of manually designed input features. On the other hand deep learning based methods can learn feature representations directly from images. Moreover, there have been proposed deep learning based method designed for MIL, e.g., (*17*). However, deep learning based method might not be directly applicable to TMB prediction problems. The main difficulties are related to computational issues caused by the nature of whole slide images, i.e., small sample size but a huge size of images. For example, TCGA bladder cancer data used in our experiments contains just hundreds of whole slide images, but either dimension of each image is at least 100,000 pixels. Each image can have hundreds or thousands image blocks even if we focus on only tumorous regions in an image. It becomes more problematic if we train a more complex deep learning model with more many parameters to be tuned. In this study, thus we pursue the traditional machine learning rather than deep learning approaches. We first calculate features from image blocks in a image and build a predictive model to aggregate the results from all the image blocks to obtain the image’s final label.

Heteroscedastic noise modeling, which assumes the noise variance to vary across the input space, provides a more realistic noise model. This idea has been widely used for Gaussian process regression (*9, 20, 21, 27*). However, in (*15*) heteroscedastic noise modeling is used for binary classification, where the noise variance is modeled as a function of privileged information that is available only during training time. The authors interpret that this latent function for the noise variance can encode the confidence about each training sample. They also have showed that this modeling improves the performance over a standard GP classifier. Our heteroscedastic modeling is quite related to this approach. We also use the heteroscedastic modeling to encode the confidence in predicting an image sample (i.e., a set of image blocks) by our prediction method in the MIL setting. However, in contrast to their method, in our method the noise variance is assumed to be a function defined on image level features that are calculated from all the image blocks in an image so that our heteroscedastic modeling is also available for test data.

## 3 Proposed Method

We propose a heteroscedastic noise model for MIL based on the framework of Gaussian process, where a latent scoring function (assumed to be follow a GP) maps from the input feature space to the intervals on the real line, and the noise variance is a latent function (assumed to be follow another GP) defined on image level features that are calculated from all the feature vectors in an image. We also propose an efficient EP algorithm to approximate the posterior distributions to learn the model hyper-parameters.

### 3.1 Data preparation

Due to computational feasibility, each whole slide image is divided into a set of image blocks. As discussed earlier, we calculate a feature vector from each image block in an image. The whole workflow of extracting image feature vectors from a whole slide image is shown in Figure 1. For detailed image processing and feature extracting steps, please refer our experimental results section (Section 4.2.1).

**Figure 1:**
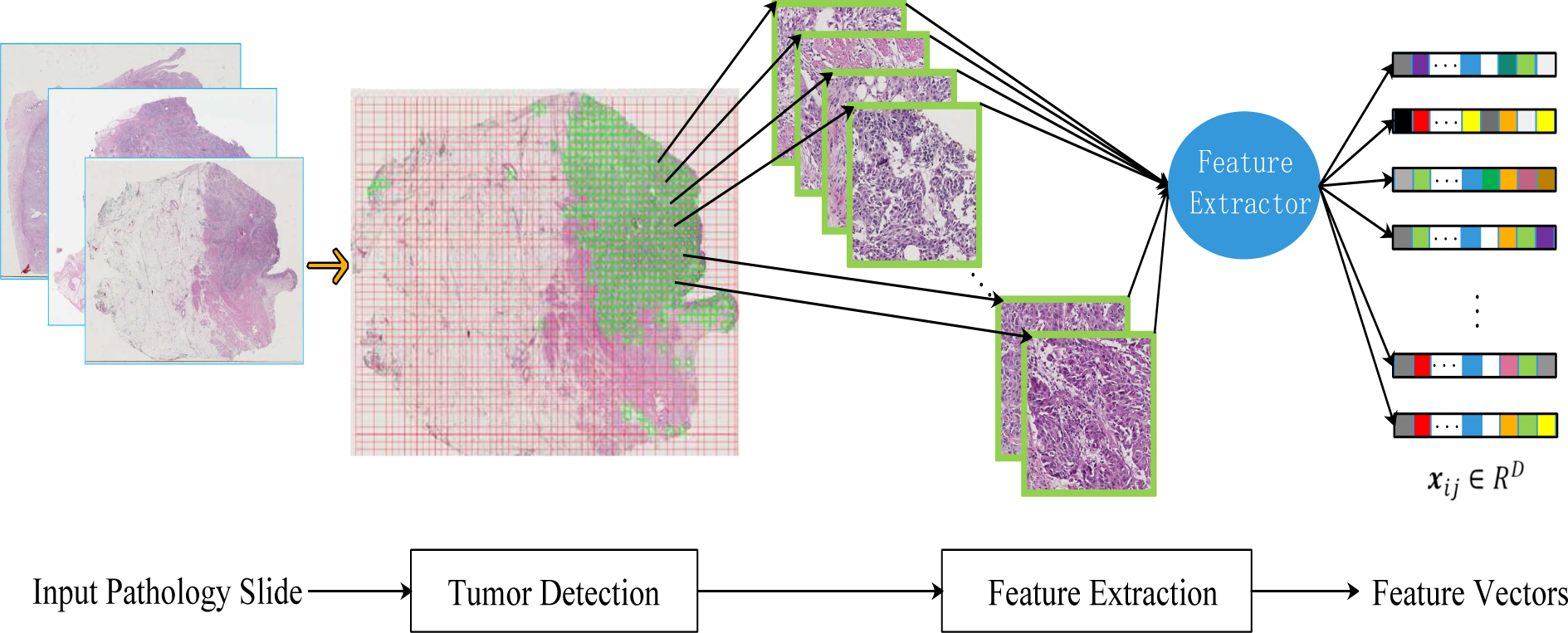
Feature extraction workflow. We divide a whole slid image into a set of image blocks and calculate a feature vector from each image block. The detailed image processing and feature extracting steps are explained in Section 4.2.1.

Let us assume that we are given a data set composed of N whole slide images. As mentioned earlier, we calculate a feature vector from each image block in an image. For example, let ***x**_ij_ ∈* ℝ^*D*^ denote as a feature vector calculated from the *j*th block in the *i*th image, and then *𝓧_i_* = {***x**_i_*_1_*, **x**_i_*_2_*, …, **x**_iN_i__*} represents a set of feature vectors of all *N_i_* blocks in the image. From the training data, we will have *𝓧*= {*𝓧*_1_*, 𝓧*_2_*, …, 𝓧_N_*} and ***y*** = [*y*_1_*, y*_2_*, …, y_N_*] ^*T*^. Here, 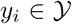 is a target label assigned to the *i*th image, where 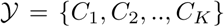 are discrete labels in the ordinal relation, i.e., *C*_1_ *≤ C*_2_ *≤ … ≤ C_K_* (*K* is set to 3 in our TBM prediction problem). Our goal is to learn a latent function that associates a set of feature vectors of an image, *𝓧_i_*, with the corresponding ordinal label *y_i_*.

### 3.2 Model specification

We first assume a latent function that maps from the input feature space to intervals on the real line, i.e., *f*: ℝ^*D*^ → ℝ and the real line is divided into non-overlapping intervals with thresholds 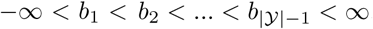. For example, the function value at ***x**_ij_* in the *i*th image, *f* (***x**_ij_*), is assumed to be in the interval, *b_y_i−1__ < t ≤ b_y_i__* (the end points, *b_y_i−1__* and *b_y_i__*, are determined by the image’s target label *y_i_*). We further assume that the function *f* (*⋅*) follows a Gaussian process:

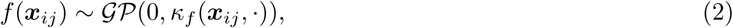

where *κ_f_* (*⋅, ⋅*) is a covariance function. In our paper, we use an automatic relevance determination (ARD) SE covariance function

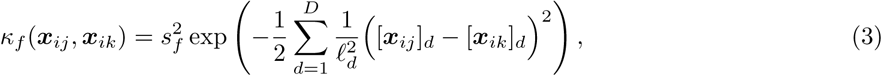

where *s_f_* is a magnitude parameter and *ℓ_d_ >* 0 is a scale length parameter which determines the relevancy of the *d*th input feature to the prediction. Letting *f_ij_* ≜ *f* (***x**_ij_*) and ***f*** be a vector containing all function values evaluated at all image blocks in training data, i.e., ***f*** = [*f*_11_*, f*_12_*, …, f_ij−_*_1_*, f_ij_, f_ij_*_+1_*, …, f_NN_i−__*_1_*f_NN_i__*], the prior distribution over the latent variable ***f*** can be given by *p*(***f***) = *𝓝*(0*, **K**_ff_*), where ***K**_ff_ ∈* ℝ^*|𝓧 |×|𝓧 |*^ = *κ_f_* (*𝓧, 𝓧*). However, since the length of ***f*** (= *|𝓧|*) can be high, we employ sparse approximation (PITC) (*22*) to reduce computational complexity. Let us define ***u** ∈* ℝ^*M*^ as inducing variables which are function values evaluated at inducing points ***Z*** ≜ {***z***_1_*, …, **z**_M_*}, where ***z**_m_ ∈* ℝ^*D*^ (*u_m_* ≜ *f*(***z**_m_*)), the conditional distribution of ***f*** given ***u*** is defined as follows

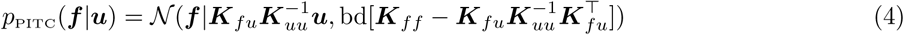

where ***K**_fu_*= *κ_f_*(*𝓧, **Z***) and ***K**_uu_* is a covariance matrix for the inducing variables, i.e., ***K**_uu_* = *κ_f_* (***Z**, **Z***) *∈* ℝ^*M ×M*^. Please note that the operator *bd*[***M***] forms a block-diagonal matrix from the matrix ***M*** where each diagonal block is calculated from each image, i.e., 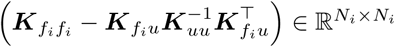, where 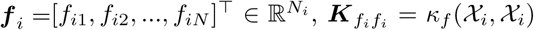 and ***K**_fu_* = *κ_f_*(*𝓧_i_, **Z***). In other words, the approximation is enabled by removing the dependency between any pairs of function values that are not in the same image.

We then propose a novel likelihood (noise) model for multiple instance learning using heteroscedastic noisy modeling, where the noise variance varies across image samples. We first need to link all the function values 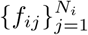 from a image to the corresponding target label *y_i_*. We will use their mean value to aggregate them into a single value, 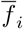, and design a likelihood function comparing that value and the corresponding target label:

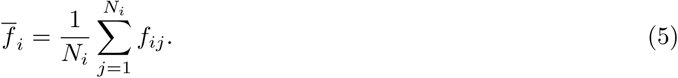

In fact, this definition can be considered as an implementation of the collective multi-instance assumption (*8, 12, 18, 32*) (where all the instances in a bag are equally contribute to the target label), rather than the standard multi-instance assumption, i.e., taking the maximum value among the instances in a bag. We hypothesis that the collective multi-instance assumption (5) is more appropriate to TBM prediction than the standard multi-instance assumption is because genetic alteration might affect the overall scanned tumor regions and thus just a single image block cannot be a dominant representative of all the others, from which we predict the TMB level of a patient sample. In addition, in contrast to the standard multi-instance assumption (using max operation), the definition (5) leads a more efficient inference algorithm which is discussed in the sequel (Section 3.3). The likelihood function could be simply defined as (please recall our definition of the latent function *f* (*⋅*)):

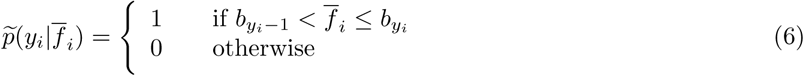

Due to noise from inputs or target labels, however, we consider a noisy version of the ideal (noise free) likelihood function (6):

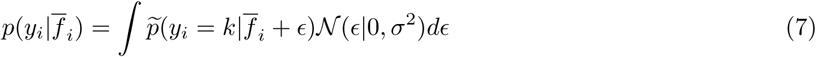

where we have assumed that the estimator 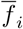 is contaminated by a Gaussian noise with zero mean and unknown variance *σ*^2^.

We further extend the likelihood function (7) to the case where the noise variance *σ*^2^ can vary across image samples, leading to a heteroscedastic noise model. We might have different levels of confidence in predicting TMB from different images. For example, we would have high confidence in prediction if the patterns across all the image blocks in an image are consistent. However, our confidence in prediction would be low if there is high diversity in the patterns across all the image blocks. Thus the noise variance in (7) can represent the level of this confidence in prediction a set of image blocks in the MIL setting. As shown in Figure 2, our main idea is to make our method put different levels of effort to classify each image correctly according to the confidence level. In other words, we make our method try to fit easier image samples well but not to put much effort in classifying harder (ambiguous) images correctly. We further assume that the noise variance can be determined from the distribution of each feature among the image blocks in an image. We define an image level feature vector 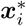 for each image, which contains statistics of each feature in an image: 

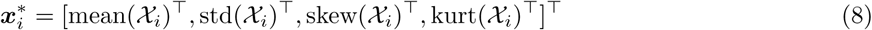

where the mean, standard deviation, skewness and kurtosis of each feature are calculated separately and thus 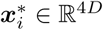. Assuming the noise variance at the *i*th image be a function of the statistical features 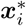, our heteroscedastic noise model can be defined as

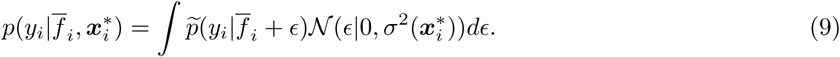

**Figure 2:**
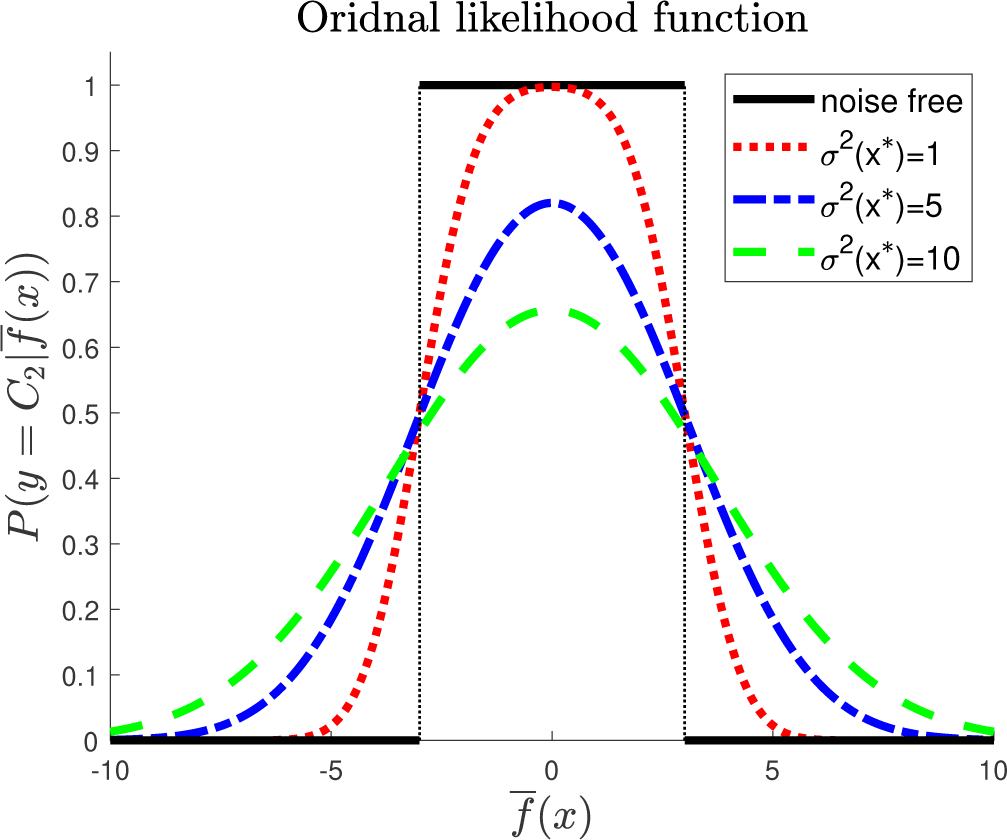
The likelihood function (9) with different levels of the noise variance. As the noise variance increases, the likelihood is reduced in overall over 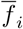. Controlling this noise variance, we can make our prediction model put different levels of effort to classify each image correctly.

Finally we assume that the noise variance 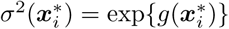 is modeled by another Gaussian process:

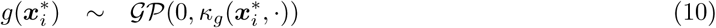

where *κ_g_* is also assumed to be a ARD SE covariance function. The statistical feature 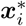 can be considered similar to the privileged information in (*15*). However, unlike the setting in (*15*), the statistical features can be always calculated from a test image.

As a summary, our predication model of TMB from images with the heteroscedastic noise modeling is given as follows:

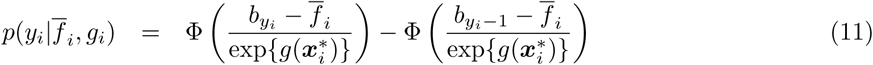

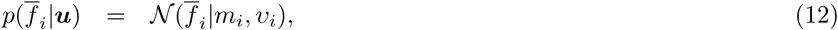

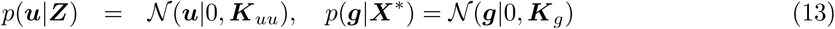

where

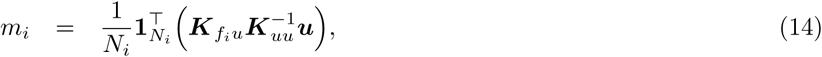

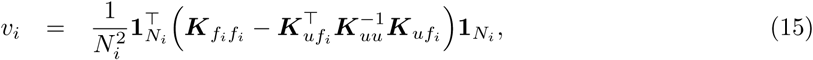

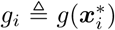 and ***g*** = [*g*_1_*, g*_2_*, …, g_N_*]^*T*^. In fact, we have included the variable 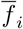 in the model for readers’ easy and clear understanding, but it can be easily marginalized out from the model.

### 3.3 Inference with EP

The inference of the model involves calculating the posterior distribution over the latent variables, ***u*** and ***g***, and estimating the hyper-parameters (e.g., scale length parameters in the ARD covariances). The posterior over ***u*** and ***g*** can be calculate as follows:

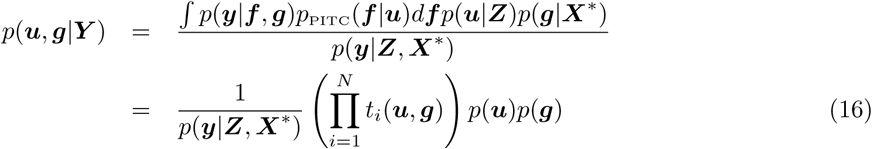

where

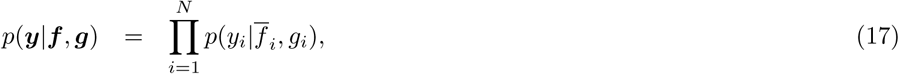

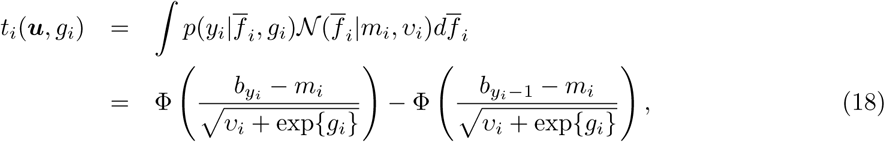

and the marginal likelihood *p*(***y**|**Z**, **X**^∗^*) (≜ *Z*) can be used to find the optimal hyper-parameters (***ϑ***) by maximizing *Z* w.r.t. ***ϑ***.

We use expectation and propagation (EP) to approximate the posterior distribution (16) as the calculations involved in (16) are intractable. EP enables analytical computation of the posterior distribution by approximating each non-Gaussian likelihood *t_i_*(***u**, g_i_*) with a non-normalized Gaussian factor 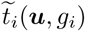, defined as follows

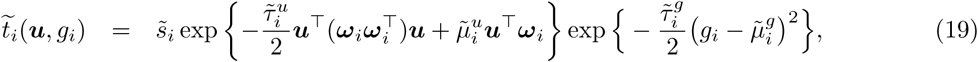

where 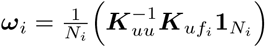. Then the approximate posterior distribution over ***u*** and ***g***, *q*(***u**, **g***), can be calculated as follows

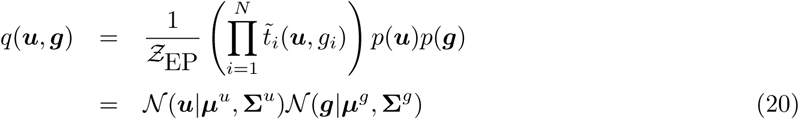

where

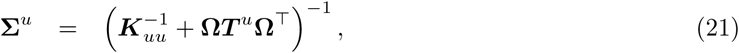

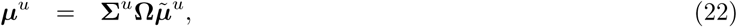

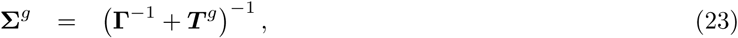

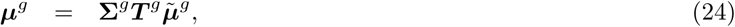

and *Ƶ*_EP_ is the EP approximation to the marginal likelihood *Ƶ*.

In the standard EP, each factor 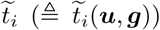 is iteratively updated until convergence in a way that it becomes the most accurate in regions of high posterior probability (*2*). The update rule is as follows. *i*) We first calculate the cavity distribution *q_\i_* by removing 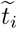 from the approximate posterior distribution *q* (≜*q*(***u**, **g***)), i.e.,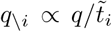. *ii*) Then we find a new distribution 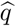 that minimizes the Kullback–Leibler divergence between 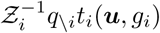 and *q*, i.e., 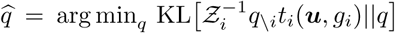, where *Z_i_* is a normalization constant. Since *q* is assumed to be Gaussian, this minimization can be done by matching the expected sufficient statistics (mean and covariance) of *q* with those of the distribution 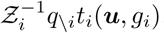. These expectation can be calculated using the derivative of log *Z_i_* w.r.t. the natural parameter of *q_\i_* (*26*). However, due to our heteroscedastic noise modeling, log *Z_i_* and its derivations cannot be calculated in closed form. However, fortunately, all the intractable calculations involve just an one-dimensional numerical integration (w.r.t. *g_i_*) which can be efficiently computed in different ways, e.g., use Gauss-Hermite quadrature. *iii*) Finally the factor 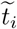 can be updated by 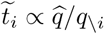. At convergence, the approximate posterior distribution can be calculated using all the factors 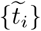 and the prior distributions as in (20). In addition, the approximate marginal likelihood also can be calculated from these factors:

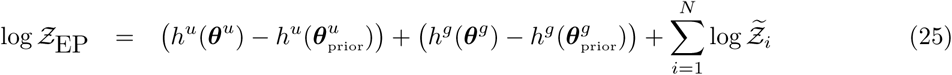

where

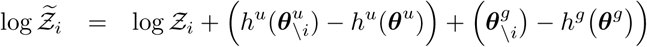

where 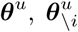 and 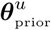 are the natural parameters of the marginal distributions, *q*, *q_\i_* and *p*(***U** |**Z***), respectively (*q* is assumed to be factorized into two independent distributions of ***u*** and ***g***, and so is *q_\i_*); and *h^u^*(***θ**^u^*) is the log partition function of a Gaussian distribution with natural parameter ***θ**^u^* (*26*). We can further optimize the hyper-parameters (***ϑ***) using the gradient of the log approximate marginal likelihood log *Z_EP_* w.r.t ***ϑ***. Please see Appendix for the detailed information about the EP algorithm for our model.

For computational efficiency, we follow the approach proposed (a variant of the standard EP algorithm) in (*14*). Instead of waiting until the EP updating converges, not only the approximate factors 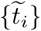 but also the hyper-parameters ***ϑ*** are updated at the same time. We use the same (batch) gradient ascent method in (*14*) to update the hyper-parameters, where each parameter has its own learning rate which is adaptively determined according to the sign change of the gradient between two consecutive iterations.

Finally, the predictive distribution of a new test image sample (e.g., a bag *𝓧*_o_ with 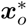) can be approximately calculated using the posterior distribution *q*(***u**, **g***) obtained from the EP algorithm:

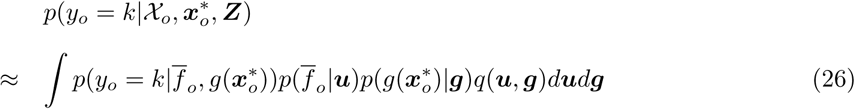

where the distribution 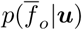 is given by

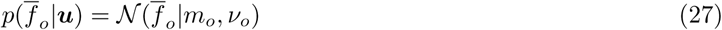

where it mean and the variance are as follows

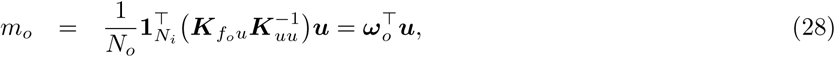

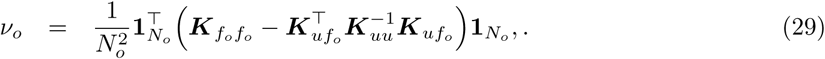

where ***K**_f_o_f_o__* = *κ_f_* (*𝓧_o_, 𝓧_o_*), ***K**_uf_o__* = *κ_f_* (***Z**, 𝓧_o_*) and **1**_*N*_o__ ∈ ℝ^*N_o_*^ is a length *N_o_* (the number of image blocks in the new image) of one vector. As a result, the predictive distribution can be calculated as

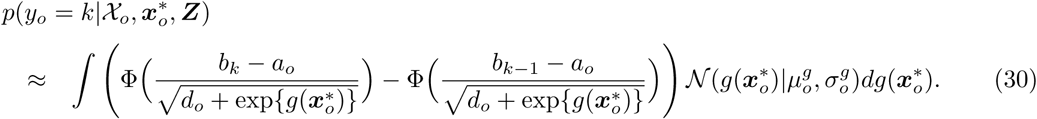

where

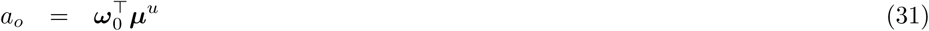

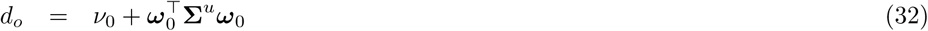

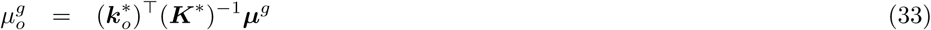

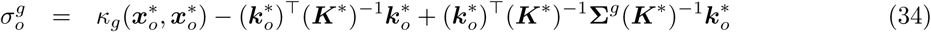

where 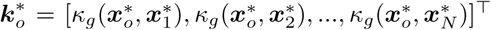. Note that this integral also does not have a close form solution. We can use again Gaussian quadrature to numerically calculate this integration.

## 4 Numerical experimental results

We investigated the performance of our prediction model on synthetic and real-world data sets. We compared our methods with the multiclass SVM method on whole slide images proposed in (*31*) (hereafter MSVM) and multiple instance ordinal regression (MIOR) (*29*). In fact, although MSVM is not a MIL method, we included this method in our experiments because the method is also based on whole slide images and the way the method combines multiple feature vectors in an image into a single vector is related to our noise modeling. MSVM also calculates the statistics of each feature, such as mean, standard deviation, skewness and kurtosis, among image blocks in an image and use these values as a new feature vector to the image which will be fed to the classifier (note that this multiclass classifier is defined by binary decomposition method, e.g., all-pair comparisons). In other words, the method uses image level features 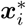 as input to the classifier. Regarding MIOR, there have been proposed very few MIL methods for ordinal regression problems. As its name suggests MIOR solves ordinal regression in the MIL setting. Note that, it is not straightforward to directly extend the standard MIL assumption to multi-class or ordinal regression problems. Thus, the method proposes its own strategy how to link multiple instances in a bag to the corresponding label. For each bag, only the instance which is nearest to the center between two corresponding hyperplanes given by the class label is selected to be compared with the label (*29*). All methods, including ours, were implemented in Matlab (version info: Matlab R2018a). For MSVM, we used LibSVM (*4*) as a base binary classifier (we only considered the linear kernel for the SVM because the nonlinear kernel, i.e., a rbf kernel, showed worse performance).

### 4.1 Synthetic data

With this synthetic data set, we tested the effectiveness of our heteroscedastic noise modeling. To do this, we considered the reduced version of our method (the Gaussian process based MIL prediction model but without the heteroscedastic noise modeling), i.e., we implemented the same Gaussian process prediction model with the likelihood function with a fixed noise variance (7), instead of the original likelihood function (9). We refer to this reduced version of our method as **GPMIL** and call our method with the heteroscedastic noise modeling by **GPMIL-HN**. Both methods were trained based on the frame of EP approach. For the case of training GPMIL, the EP algorithm becomes more simplified because it involves neither the additional Gaussian process which models the noise variance from image level features 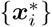 nor any numerical integration, e.g., Gauss-Hermite quadrature, as in the case of learning GPMIL-HN.

To evaluate the performance of the methods, we calculated the multi-class prediction accuracy, which is given by

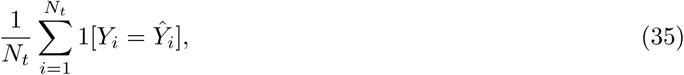

where *N_t_* is the number of the total test samples, *Ŷ*_*i*_ is the estimated label for the *i*th sample and the operator 1[expr] is 1 if the expression, expr, is true or 0 otherwise. To compare the performance of the methods, we used random K-fold cross-validations where the data points (bags in our case) are randomly divided into K-folds and 1 fold is used to test the prediction performance of the model and the rest folds are used to train the model. For MSVM and MIOR, the training data (bags in the *K −* 1 folds) are again randomly divided into training and validation sets to select the optimal hyper-parameters, e.g., the regularization constant. However, for our methods all the training data can be used to train the model and to optimize the hyper-parameters, such as ARD coefficients and inducing points, because our methods provide approximate marginal likelihood from which we can optimize these hyper-parameters.

#### 4.1.1 Dataset

We generated a 2 dimensional three class MIL data set, where each point (instances) in a sample (bag) is generated from one of three Gaussian distributions or the mixture of them. Please see Figure 3, where a square with different colors represents the mean of each Gaussian distribution (mean: *C*_1_ = [*−*3*, −*3]^*T*^, *C*_2_ = [*−*0.5*, −*3]^*T*^, *C*_3_ = [3, 3]^*T*^ and covariance matrix: *C*_1_ = [1, 0.2; 0.2, 1], *C*_2_ = [1, 0.5; 0.5, 1], *C*_3_ = [1, 0; 0, 1]). Note that these mean vectors and covariance matrices were chosen arbitrarily. We generated 100 bags from each class and made each bag to have 100 instances. We assumed that there were two types of samples: easy and hard samples. For the case of easy samples, all instances in a bag were sampled from a single Gaussian distribution corresponding to the label to which the bag belongs. On the other hand, for the case of hard samples, instances in a bag were sampled from a mixture of the three Gaussian distributions with the mixing coefficients [0.4, 0.3, 0.3]. We made sure that the mixing coefficient for the corresponding Gaussian distribution to the class to which a bag belongs was always the largest, i.g., it was set to 0.4, so that the majority of instances in a bag still came from the corresponding class. Finally we introduce the noise rate which is the portion of hard sample in the total samples, i.e., if the noise rate is set to 0.5, then it means that 50 bags among 100 bags in each class are hard samples.

**Figure 3:**
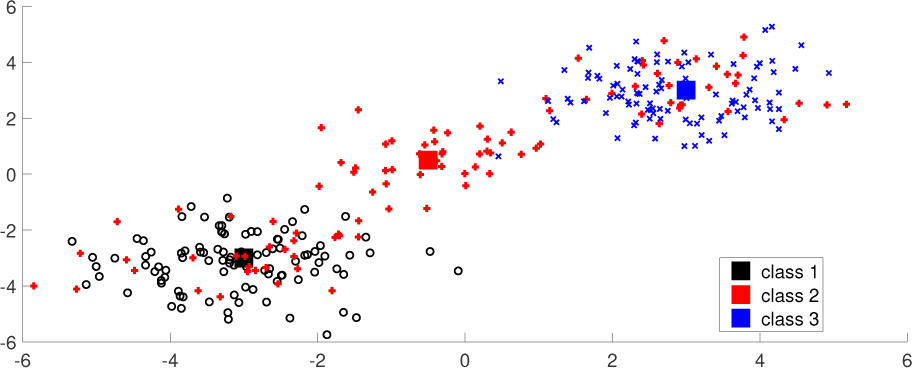
2-D 3 class synthetic data. Each square in different colors represents the mean vector of each Gaussian distribution (each class) and the symbols, (x, o, +) are instances in class 1, 2 and 3, respectively. Here, we show just three samples (bags) from each class as examples: i.e., two easy samples, black (class 1) and blue (class 3), and one hard sample (red). The instances in the red bag are spread across the areas where samples from the other classes mainly reside.

**Figure 4:**
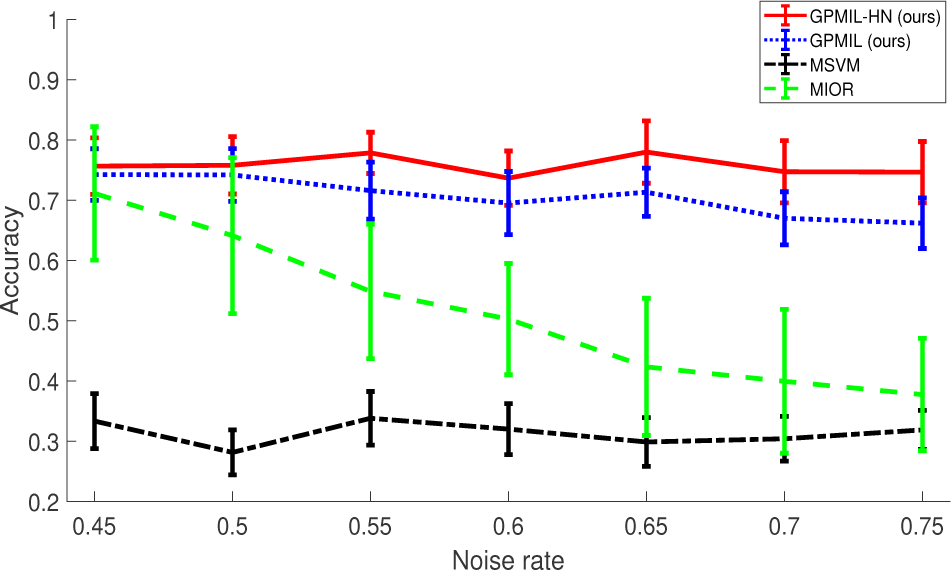
The accuracy of each method evaluated with increasing the noise rate. Each line and error bar represent the mean performance and the corresponding standard deviation, respectively, of each method evaluated by 5-repetition of random 5-fold cross validation.

#### 4.1.2 Performance evaluation

We evaluated the performance of the methods with increasing the noise rate. We repeated random 5-fold cross validation 5 times at each noise rate and report the results in Figure (4). Each line represents the mean performance of each method and each error bar is the standard deviation calculated from the repetition at each noise level. We first note from the figure that MSVM is not able to handle the presence of noise in the MIL setting. Its performance is almost the same as that of a random classifier. The figure also showed that MIOR worked better than MSVM but its performance decreased as the noise rate increased. It might indicate that MIOR is sensitive to noise. It might be because the method selects only a single instance from each bag to train the model. It is more likely that an irrelevant instance to a bag can be selected as a training example in the presence of highly noisy data. However, our methods, both GPMIL-HL and GPMIL, showed almost stable performance across all the range of the noise rates. It might be because our methods are based on the collective MIL assumption which use all instances in a bag to train the model, unlike the assumption used in MIOR or the standard MIL assumption. We also confirmed that there is a gain caused by our heteroscedastic noise modeling from the comparison between GPMIL-HN and GPMIL (without the heteroscedastic noise modeling).

### 4.2 TMB prediction

#### 4.2.1 Dataset

A cohort of 386 bladder cancer patients (and corresponding clinical information) with 457 diagnostic H&E stained whole slide images were downloaded from TCGA data portal. Among 386 patient samples, 24 patients were excluded due to poor quality of their images, such as undesired pen marks, torn tissues or other artifacts. According to the percentile of total number single nucleotide variants (*24*), the remaining 362 patient samples had been categorized into 3 groups: 119 low TMB patients, 123 medium TMB patients and 120 high TMB patients. Although a small number of patients includes more than 1 diagnostic slides, for simplicity this study only uses the first diagnostic slide (i.e., with DX1 suffix) of those patients. Therefore, the dataset for evaluating TMB prediction consists of 362 different patients of whole slides images, with each image scanned at 20*×* or 40*×* magnification and having a size from hundreds of megabytes to several gigabytes.

#### 4.2.2 Image processing pipeline

Since the whole slide image has a large volume size and is difficult to be processed directly, we use a multi-resolution analysis pipeline to extract features from the image. Fig. illustrates our image processing pipeline, which includes two main modules: tumor detection and feature extraction. In the tumor detection module, the whole slide image is processed at a coarse level (e.g., 5*×* in this study). First, the whole slide image is divided into a number of non-overlapping 256*×*256 image blocks (see overlapped red and green blocks in the image). Next, local binary patterns (LBP) texture features are computed from each image block. The blocks that belong to tumor regions (see green blocks in the image) are then detected by using a trained SVM classifier (*30*). In the feature extraction module, the detected tumor blocks are analyzed at a fine level (e.g., 10*×* in this study). To extract fine level features, we make use of transfer learning on the Resnet50 model (*13*), which was pre-trained on approximately 1.28 million image from ImageNet dataset. We replace the output layer of Resnet50 model with a global average pooling layer, and hence the Resnet50 model finally outputs a 2048 dimensional feature vector for each input tumor block. Finally the dimensional of input features was further reduced to 50 by principal component analysis (PCA) since these first 50 principal components explain 97% of variance.

#### 4.2.3 Performance evaluation

We evaluated the performance of each method again with random 5-fold cross-validation. We repeated each experiment 5 times. Beside the prediction accuracy, we computed F_1_ measure, which is the Harmonic mean of precision and recall, i.e., 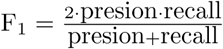 of each class under the assumption that the class is supposed to be the positive class and the other classes negative. Calculating F_1_ measures, we can examine the performance of each method with respect to a specific class.

Table 1 shows the multi-class prediction accuracy of each method. Although, the overall performance of all the method is not that impressive, comparing GPMIL-HN and GPMIL, we can see a clear improvement due to the heteroscedastic noise modeling. In fact, GPMIL did not work at all for the TMB prediction as the method classified all images as *intermediate* class as shown in Table 2. Our method also showed better performance than MSVM and MIOR. In particular, MIOR also did not work for the TMB prediction. It classified most all test images as the same class 20 times during the experiments, where the total number of evaluations was 25 (= 5 repetition *×* 5 folds). Please see the results of MIOR in Table 2. As a result, we confirmed from the TMB prediction on the real-world data set (TCGA bladder cancer whole slide images) that the usefulness of our heteroscedastic noise modeling.

**Table 1:** The multi-class prediction accuracy. We report the mean performance of each method. The number in the parentheses represents the standard deviation.

| Method | Accuracy |
| --- | --- |
| GPMIL-HN(ours) | **0.429** (0.055) |
| GPMIL (ours) | 0.339 (0.018) |
| MSVM | 0.346 (0.053) |
| MIOR | 0.341 (0.022) |

**Table 2:** The F1 measures. We report the mean performance of each method and its standard deviation in the parentheses. The symbol ’-’ means that the value is non-available. This is happen when the method does not classify any test image samples as the corresponding class in any-fold experiment during 5-fold cross-validation (5 repetition).

| Method | (TMB) Low | Intermediate | High |
| --- | --- | --- | --- |
| GPMIL-HN (ours) | 0.429 (0.084) | 0.466 (0.044) | 0.364 (0.097) |
| GPMIL (ours) | - | 0.504 (0.017) | - |
| MSVM | 0.196 (0.098) | 0.391 (0.062) | 0.405 (0.090) |
| MIOR | - | - | - |

## 5 Conclusion

In this study we develop a computational prediction method of the TMB level of patients from whole slide images, where the prediction problem is formulated as a MIL problem and the heteroscedastic noise model is used to model the noise variance from image level features. In particular, this noise variance encodes the confidence that our method has regarding image samples. The method trusts image samples that are easier (having the higher confidence level or the lower noise variance) but less relies on image samples that are hard (having the lower confidence level or the higher noise variance). Then the posterior distributions and the model hyper-parameters are efficiently learned by the EP algorithm. We finally have demonstrated from the synthetic and TCGA bladder cancer whole slide image data sets the usefulness of our heteroscedastic noise modeling over the other baseline methods.

In comparison with MIOR, which is also an ordinal regression method in the MIL setting, our method has several advantages. As we have seen from the experimental results on the synthetic data, MIOR is more sensitive to noisy instances presented in a bag than our method is. It would be mainly because of its selective assumption, i.e., only the instance which is nearest to the center between two hyperplanes given by the class label is selected to train the model (*29*). In addition, our method is a Bayesian approach and thus would be more robust against overfitting than MIOR (which is implemented by regularized loss minimization) is. Furthermore, the marginal likelihood allows us to use more complex covariance functions, corresponding to the kernel function used in in MIOR. With using of ARD covariance functions we can automatically reduce the influence of features that are irrelevant to the prediction task. We also can optimize the parameters of in ARD covariance functions. This advanced adjustment is not available with MIOR.

## Appendix

### Experiment setting

The hyper-parameters in our method include inducing points ***Z*** and model-parameters for the covariance functions, *κ_f_* and *κ_g_*. In all our experiments, the number of inducing points, *M*, was set to 100. Each inducing point was optimized by maximizing the marginal likelihood in the framework of EP. They were initialized by randomly selected feature vectors from the training data. In addition, we set *b*_1_ and *b*_2_ to 0 and 2 respectively. In fact, these values also can be optimized. However, we did not find significant improvement from optimizing of these values. For MSVM, the regularized constant was selected among [2^*−*3^; 2^*−*2^; 2^*−*1^; 2^0^; 2^1^; 2^2^; 2^3^; 10; 2^4^] by cross-validation. For MIOR there are two regularized constants. These values were also chosen by CV. The search space was 2-dimensional grid space and the possible candidates for each parameter were set to the same set, [2^*−*2^; 2^*−*1^; 2^0^; 2^1^; 2^2^; 2^3^].

### Detailed derivations

The detailed steps for the EP in Section 3.3 are as follows.

*i)* The cavity distribution 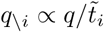 can be calculated as

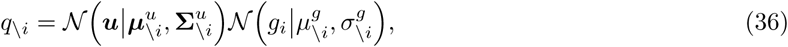

where

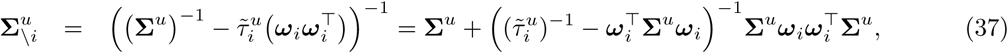

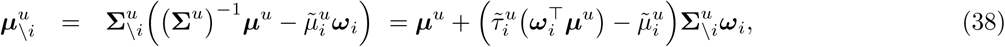

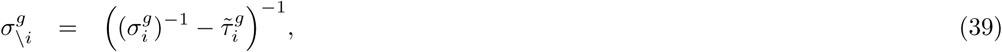

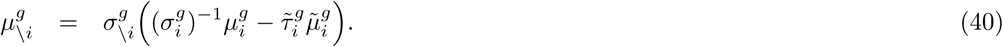

ii) We now compute the distribution 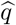:

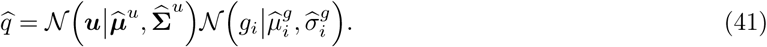

To do this, we first need to calculate *Z_i_*, i.e., the normalization constant of *t*(***u**, g_i_*)*q_\i_*,:

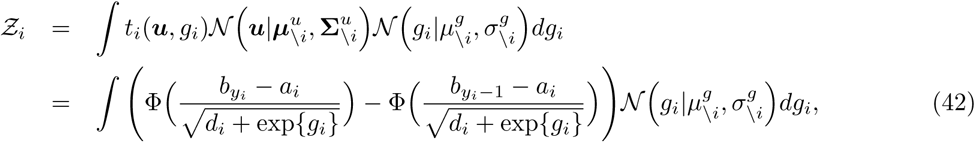

where

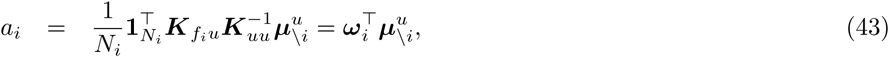

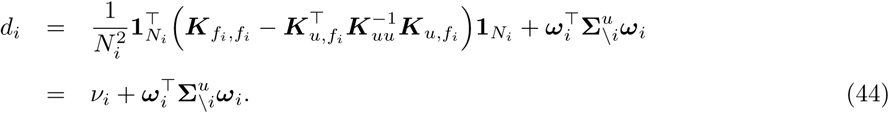

Note that (42) does not admit a close-form solution and thus we need numerical approximation methods, e.g., Gauss-Hermite quadrature. Then, the distribution 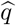 can be computed from the derivative of log *Ƶ_i_* with respect to the parameters of *q_\i_*. The mean 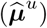 and covariance matrix 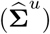 of the distribution 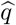 is given by

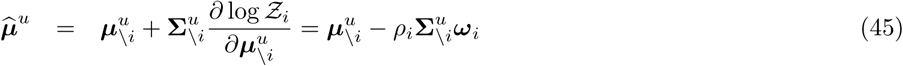

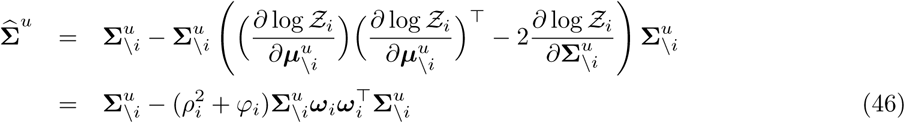

where

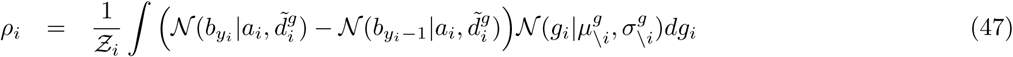

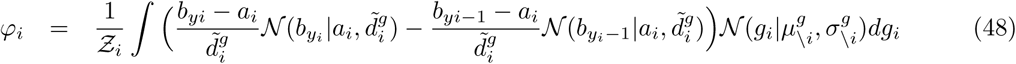

where 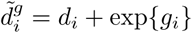. Then 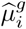 and 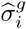 are computed by

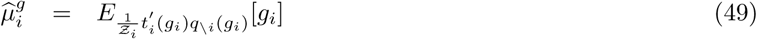

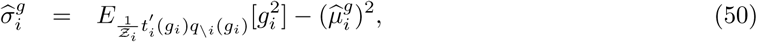

Where 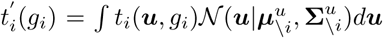 and these computations can be done by numerical methods, e.g., Gauss-Hermite quadrature.

*iii)* The factor 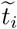 can be updated by 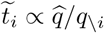.

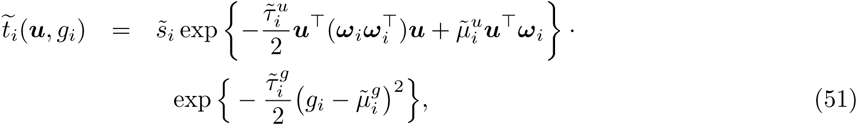

For the latent variable ***u***, the updated factor is given by

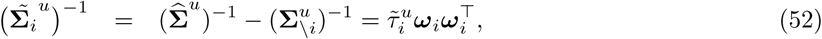

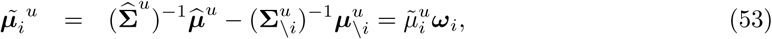

where

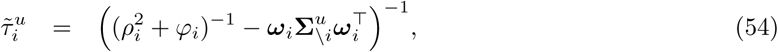

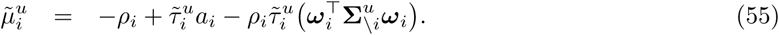

For the latent variable *g_i_*, the updated factor is given by

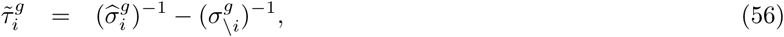

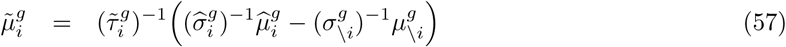

The approximate marginal likelihood *Ƶ*_EP_ = *p*(***y**|**Z**, **X**^∗^*) can be calculated as

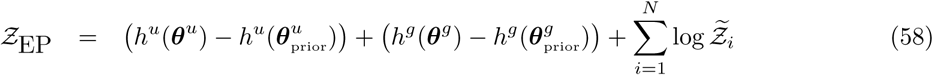

where

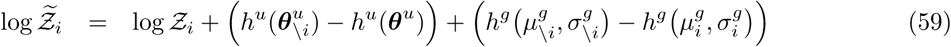

With the definition of the log-normalizer of a Gaussian distribution, *Z*_EP_ can be computed as

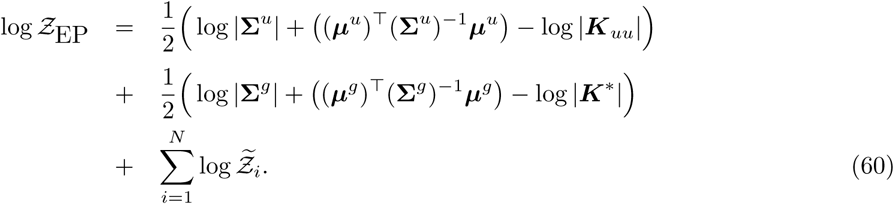

The gradient of *Z*_EP_ with respect to a hyper parameter, *ϑ*, in the covariance function *κ_f_* can be computed as

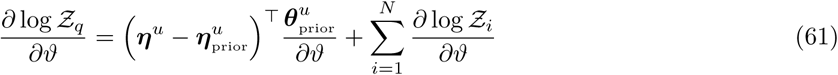

where

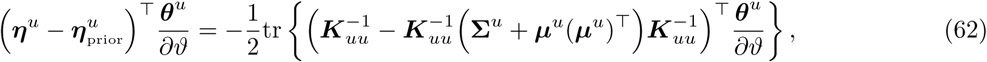

and

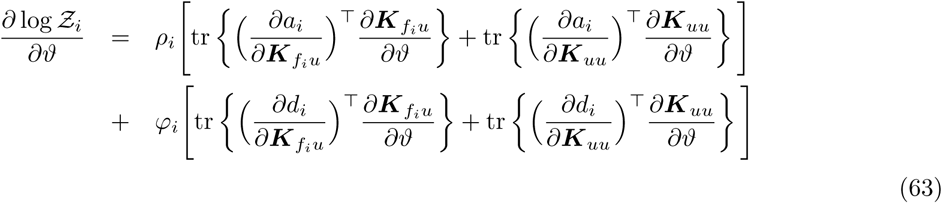

The gradient of *Ƶ*_EP_ with respect to a hyper parameter in the covariance function *κ_g_*, can be further simplified (since 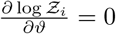, ∀i in this case):

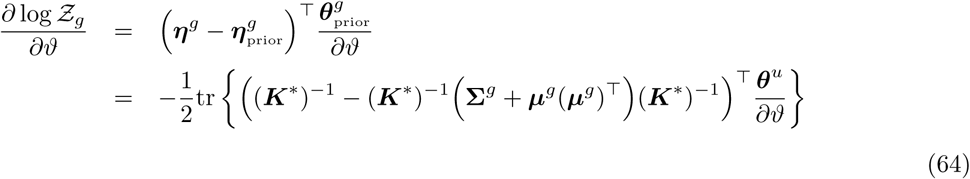

## References and Notes

1. Stuart Andrews, Ioannis Tsochantaridis, and Thomas Hofmann. Support vector machines for multiple-instance learning. In Proceedings of the 15th International Conference on Neural Information Processing Systems, Advances in Neural Information Processing Systems (NIPS), pages 577–584, 2002.

2. Christopher M. Bishop. Pattern Recognition and Machine Learning (Information Science and Statistics). Springer-Verlag, Berlin, Heidelberg, 2006.

3. R. L. Brown, S. D. nad Warren, E. A. Gibb, S. D. Martin, J. J. Spinelli, B. H. Nelson, and R. A. Holt. Neo-antigens predicted by tumor genome meta-analysis correlate with increased patient survival. Genome research, 25:743–50, 2014.

4. C. C. Chang and C. J. Lin. LIBSVM: A library for support vector machines, 2001.

5. Wei Chu and Zoubin Ghahramani. Gaussian processes for ordinal regression. Journal of Machine Learning Research, 6:1019.

6. Nicolas Coudray, Paolo Santiago Ocampo, Theodore Sakellaropoulos, Navneet Narula, Matija Snuderl, David Fenyö, Andre L. Moreira, and Narges Razavian.

7. Thomas G. Dietterich, Richard H. Lathrop, and Tomás Lozano-Pérez. Solving the multiple instance problem with axis-parallel rectangles. Artificial Intelligence, 89(1):31–71, 1997.

8. James Foulds and Eibe Frank. A review of multi-instance learning assumptions. The Knowledge Engineering Review, 25(1):1–25, 2010.

9. Paul W. Goldberg, Christopher K. I. Williams, and Christopher Bishop. Regression with input-dependent noise: A gaussian process treatment, 1998.

10. Aaron M. Goodman, Shumei Kato, Lyudmila Bazhenova, Sandip P. Patel, Garrett M. Frampton, Vincent Miller, Philip J. Stephens, Gregory A. Daniels, and Razelle Kurzrock. Tumor mutational burden as an independent predictor of response to immunotherapy in diverse cancers. Molecular Cancer Therapeutics, 16(11):2598–2608, 2017.

11. M. Haußmann, F. A. Hamprecht, and M. Kandemir. Variational bayesian multiple instance learning with gaussian processes. In Proceedings of the IEEE International Conference on Computer Vision and Pattern Recognition (CVPR), pages 810–819, 2017.

12. Jianjun He, Hong Gu, and Zhelong Wang. Bayesian multi-instance multi-label learning using gaussian process prior. Machine Learning, 88(1):273–295, Jul 2012.

13. Kaiming He, Xiangyu Zhang, Shaoqing Ren, and Jian Sun. Deep residual learning for image recognition. In Proceedings of the IEEE conference on computer vision and pattern recognition, pages 770–778, 2016.

14. Daniel Hernandez-Lobato and Jose Miguel Hernandez-Lobato. Scalable gaussian process classification via expectation propagation. In Proceedings of the International Conference on Artificial Intelligence and Statistics (AISTATS), volume 51 of Proceedings of Machine Learning Research, pages 168–176, Cadiz, Spain, May 2016. PMLR.

15. Daniel Hernández-lobato, Viktoriia Sharmanska, Kristian Kersting, Christoph H Lampert, and Novi Quadrianto. Mind the nuisance: Gaussian process classification using privileged noise. In Advances in Neural Information Processing Systems 27.

16. Le Hou, Dimitris Samaras, Tahsin M. Kurç, Yi Gao, James E. Davis, and Joel H. Saltz. Patch-based convolutional neural network for whole slide tissue image classification. Proceedings of the IEEE International Conference on Computer Vision and Pattern Recognition (CVPR), pages 2424–2433, 2016.

17. Maximilian Ilse, Jakub M. Tomczak, and Max Welling. Attention-based deep multiple instance learning. In Proceedings of the International Conference on Machine Learning (ICML), 2018.

18. Melih Kandemir, Manuel Haussmann, Ferran Diego, Kumar Rajamani, Jeroen Van Der Laak, and Fred Hamprecht. Variational weakly supervised gaussian processes. In British Machine Vision Conference, York, UK, 2016.

19. Minyoung Kim and Fernando Torre. Multiple instance learning via gaussian processes. Data Mining and Knowledge Discovery, 28(4):1078–1106, 2014.

20. Miguel Lázaro-Gredilla and Michalis K. Titsias. Variational heteroscedastic gaussian process regression. In Proceedings of the International Conference on Machine Learning (ICML), pages 841–848, 2011.

21. N. Quadrianto, K. Kersting, M. D. Reid, T. S. Caetano, and W. L. Buntine. Kernel conditional quantile estimation via reduction revisited. In Proceedings of the IEEE International Conference on Data Mining (ICDM), pages 938–943, 2009.

22. Joaquin Quiñonero Candela and Carl Edward Rasmussen. A unifying view of sparse approximate gaussian process regression. J. Mach. Learn. Res., 6:1939–1959, December 2005.

23. Carl Edward Rasmussen and Christopher K. I. Williams. Gaussian Processes for Machine Learning (Adaptive Computation and Machine Learning). The MIT Press, 2005.

24. A Gordon Robertson, Jaegil Kim, Hikmat Al-Ahmadie, Joaquim Bellmunt, Guangwu Guo, Andrew D Cherniack, Toshinori Hinoue, Peter W Laird, Katherine A Hoadley, Rehan Akbani, et al. Comprehensive molecular characterization of muscle-invasive bladder cancer. Cell, 171(3):540–556, 2017.

25. Andrew J Schaumberg, Mark A Rubin, and Thomas J Fuchs. H&e-stained whole slide image deep learning predicts spop mutation state in prostate cancer. bioRxiv, 2018.

26. Matthias Seeger. Expectation propagation for exponential families. Technical report, 2005.

27. Ville Tolvanen, Pasi Jylänki, and Aki Vehtari. Expectation propagation for nonstationary heteroscedastic gaussian process regression. 2014 IEEE International Workshop on Machine Learning for Signal Processing (MLSP), pages 1–6, 2014.

28. Nils Weidmann, Eibe Frank, and Bernhard Pfahringer. A two-level learning method for generalized multi-instance problems. In Proceedings of the European Conference on Machine Learning (ECML), pages 468–479, Berlin, Heidelberg, 2003.

29. Y. Xiao, B. Liu, and Z. Hao. Multiple-instance ordinal regression. IEEE Transactions on Neural Networks and Learning Systems, 29(9):4398–4413, 2018.

30. Hongming Xu, Cheng Lu, Richard Berendt, Naresh Jha, and Mrinal Mandal. Automated analysis and classification of melanocytic tumor on skin whole slide images. Computerized Medical Imaging and Graphics, 66:124–134, 2018.

31. Hongming Xu, Sunho Park, and Tae Hyun Hwang. Automatic classification of prostate cancer gleason scores from digitized whole slide tissue biopsies. In 17th International Workshop on Data Mining in Bioinformatics, 2018.

32. Xin Xu. Statistical learning in multiple instance problems. Master’s thesis, University of Waikato, Hamilton, New Zealand, 2003.

